# Epigenetic CRISPR Screening of 9p21.3 Non-Coding Regions identifies Cis-Regulatory Elements of P16^INK4a^ and P15^INK4b^ Controlling Cellular Senescence

**DOI:** 10.1101/2025.09.29.679215

**Authors:** Jiping Yang, HyeRim Han, Xifan Wang, Yousin Suh

## Abstract

Cellular senescence is a hallmark of aging and a promising target for extending human healthspan. Senescence is often accompanied by upregulation of the key senescence marker gene *CDKN2A*, yet the mechanism underlying its transcriptional activation remains unclear due to complex *cis*-regulations within the 9p21.3 locus. Here, we performed complementary CRISPR activation and interference screens in human mesenchymal stromal cells (MSCs) to systematically map non-coding *cis*-regulatory elements (CREs) at this locus that epigenetically regulate senescence. This approach revealed senescence-regulating CREs (SenReg-CREs) that bidirectionally modulate senescence through P16^INK4a^ and P15^INK4b^. Notably, we identified a primate-specific short interspersed nuclear element (SINE) MIR3 embedded within the most potent distal SenReg-CRE. Deletion of this SINE:MIR3 accelerated senescence, revealing its potential insulator function in restraining *CDKN2A/CDKN2B* activation. Collectively, these findings reveal novel mechanisms underlying senescence-associated transcriptional activation of *CDKN2A/CDKN2B* and demonstrate that senescence is malleable through manipulation of regulatory element activity, highlighting the potential of epigenetically targeting these SenReg-CREs for senomorphic interventions.

## Introduction

Cellular senescence is a hallmark of aging^1^, characterized by irreversible cell cycle arrest in response to stress or damage. Accumulation of senescent cells contributes to age-related tissue dysfunction and diseases^2-5^. The cyclin-dependent kinase inhibitor p16^INK4a^ is a widely used biomarker of cellular senescence, typically low in proliferating cells but strongly upregulated during senescence^6-8^. *In vitro* suppression of P16^INK4a^ delays cellular senescence^*9*^. I*n vivo* clearance of p16^Ink4a^-expressing cells preserves tissue function, delays the onset of age-related pathologies, and extends healthspan and lifespan in both progeroid and naturally aged mice^4,10^, establishing p16^INK4a^ as a compelling target for senescence modulation.

P16^INK4a^ is encoded by the *CDKN2A* gene, which also produces P14^ARF^ through an alternative transcript. Adjacent to *CDKN2A* is *CDKN2B*, which encodes another cyclin-dependent kinase inhibitor P15^INK4b^, another gene frequently upregulated during cellular senescence. Both genes are located within the 9p21.3 locus, notable for its large gene desert. Previous studies have revealed that this gene desert contains multiple cis-regulatory elements (CREs) that play critical roles in controlling *CDKN2A* and *CDKN2B* expression^11,12^. Nevertheless, how changes in these CREs’ activity regulate cellular senescence remains poorly understood. Recent advances in CRISPR screening approaches that facilitate direct manipulation of CRE activity^13-16^ enable comprehensive mapping of functional CREs at 9p21.3 and elucidation of their roles in cellular senescence.

Mesenchymal stromal cells (MSCs) hold therapeutic potential for age-related conditions^17^. However, their therapeutic efficacy is compromised by cellular senescence that accumulates during *in vitro* expansion and in *in vivo* niche^18,19^. Genetic enhancement strategies have successfully generated senescence-resistant MSCs and enhanced stem cell functions^20-22^, but these approaches require coding sequence modifications, raising safety concerns. In contrast, epigenetic modulation by targeting non-coding regulatory elements offers a potentially safer strategy to counteract MSC senescence without altering protein-coding sequences. Leveraging this potential, however, requires a deeper dissection of the *cis-*regulatory landscape governing cellular senescence, including the CREs orchestrating senescence marker genes *CDKN2A/CDKN2B*.

In this study, we hypothesized that non-coding CREs at 9p21.3 can regulate P16^INK4a^ and be epigenetically targeted to counteract senescence in MSCs. To test this, we first mapped CREs activated during cellular senescence within a 1.25Mb non-coding genomic region of 9p21.3 and performed CRISPR activation and interference screens in a human MSC senescence model. These screens revealed two senescence-regulating CREs (SenReg-CREs) with bidirectional effects: activation accelerates senescence, whereas inhibition delays senescence, with P16^INK4a^ and P15^INK4b^ as their target genes. Notably, we discovered a primate-specific short interspersed nuclear element (SINE) MIR3 embedded within the most potent SenReg-CRE, which appear to function as an insulator, as its deletion accelerates senescence. Collectively, our work uncovers a new mechanism underlying *CDKN2A/CDKN2B* activation during cellular senescence and provides the first proof-of-concept that epigenetic modulation of distal non-coding SenReg-CREs can slow down cellular aging, offering a novel strategy to enhance cellular resilience by targeting non-coding regulatory elements.

## Results

### Mapping *cis*-regulatory elements activated during MSC senescence

We employed a replicative senescence model using human MSCs differentiated from human embryonic stem cells (ESCs). MSCs entered a growth arrest state by passage 20 **(Fig. S1A)**. Senescent MSCs (SEN MSCs) exhibited an enlarged and flattened morphology **(Fig. S1A)**, along with a marked increase in the proportion of senescence-associated β-gal (SA-β-gal) positive cells **(Fig. S1B)** compared to proliferating MSCs (PRO MSCs). Senescence-associated molecular changes including upregulation of P16^INK4a^, P21^CIP1^ and IL6, along with a reduction in nuclear lamina gene Lamin B1, were observed in SEN MSCs **(Fig. S1C)**, confirming the successful establishment of cellular senescence.

To assess CRE activity, we performed ATAC-seq on PRO (passage 3) and SEN (passage 17 and passage 18). We focused on ATAC-seq peaks uniquely detected in SEN MSCs but absent in PRO MSCs (SEN MSC-specific CREs, SEN-CREs) **(Fig. S1D)**, reflecting their increased chromatin accessibility during MSC senescence. To gain additional insights into their epigenetic states, we applied the 18-state ChromHMM model^23^ for chromatin annotation. Interestingly, these SEN-CREs were enriched in weak enhancers and quiescent regions **(Fig. S1E)**. Given that the 18-state ChromHMM model was generated from proliferating ESC-differentiated MSCs, it is likely that senescence led to the enhanced activation of weak enhancers and opening of previously quiescent regions.

### Expression changes of *CDKN2A* and *CDKN2B* during MSC senescence

Next, we focused on the 9p21.3 locus harboring *CDKN2A* and *CDKN2B*. In addition to P14^ARF^ and P16^INK4a^, *CDKN2A* encodes three other transcript variants including P16^gamma^, P12 and isoform 6 **(Fig. 1A)**. To determine which transcript variants were expressed in MSCs and how their expression changes during senescence, we analyzed the transcriptome of PRO and SEN MSCs. As expected, the expression of first exon of P16^INK4a^ was upregulated whereas the first exon of P14^ARF^ showed no significant change during senescence **(Fig. 1A)**, indicating that senescence is associated with specific upregulation of P16^INK4a^. The other transcript variants were not detectable **(Fig. 1A)**, as their unique exons showed no expression, suggesting they are not expressed in MSCs. Additionally, P15^INK4b^ was confirmed to be upregulated during senescence **(Fig. 1A)**, although expression difference between its two isoforms could not be distinguished.

**Figure 1.**
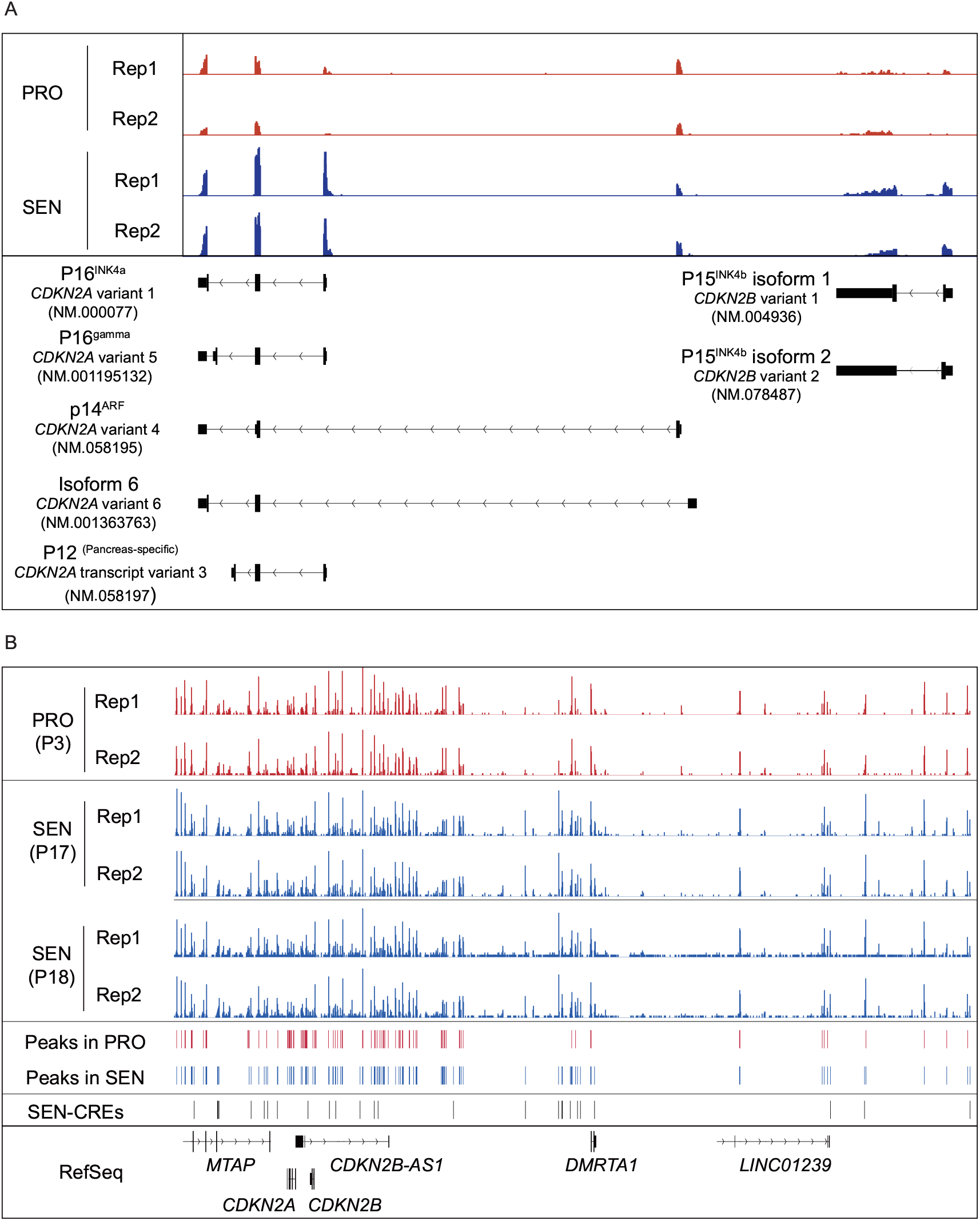
Identification of Senescence-specific Cis-Regulatory Elements (SEN-CREs) at the 9p21.3 Locus. **(A)** RNA-sequencing tracks for proliferating (PRO) and senescent (SEN) mesenchymal stromal cells (MSCs) at *CDKN2A and CDKN2B*. Increased expression of senescence-associated transcripts, including *P16*^*INK4a*^ and *P15*^*INK4b*^, is observed in SEN MSCs. **(B)** Chromatin accessibility profiles in PRO (passage 3) and SEN (passages 17 and 18) MSCs at *9p21*.*3* locus. Tracks show chromatin peaks called in PRO MSCs, SEN MSCs, and identified 9p21.3 SEN-CREs. The genomic locations of key genes *MTAP, CDKN2A, CDKN2B, CDKN2B-AS1, DMRTA1*, and *LINC01239* are indicated.

### Selection of candidate 9p21.3 SEN-CREs regulating *CDKN2A/CDKN2B* for CRISPR screens

The gene desert region upstream of *CDKN2B*, which overlaps with the long non-coding RNA gene *CDKN2B-AS1*^11,24^, can regulate the expression of P16^INK4a^ and P15^INK4b^, strongly suggesting that this non-coding region harbors CREs for *CDKN2A/CDKN2B*. However, whether the distal upstream, intergenic, and downstream regions of *CDKN2A/CDKN2B* also function as CREs and their roles in the context of cellular senescence remain unclear. We hypothesized that the increased activity of SEN-CREs at 9p21.3 upregulates P16^INK4a^ and P15^INK4b^, therefore contributing to cellular senescence. To investigate this, we selected 25 candidate Sen-CREs **(Fig. 1B)** within a 1.25Mb region spanning protein-coding genes *MTAP, CDKN2A, CDKN2B, DMRTA1*, and long-noncoding RNA genes *CDKN2B-AS1* and *LINC01239* for CRISPR screens.

### CRISPRi screen reveals 9p21.3 SEN-CREs capable of decelerating senescence

To identify functional SEN-CREs - regulatory elements activated during senescence and whose inhibition can delay the process, we performed a CRISPR - interference (CRISPRi) screen targeting candidate 9p21.3 SEN-CREs in a long-term MSC replicative senescence model **(Fig. 2A)**. CRISPR sgRNA libraries were designed to achieve saturated coverage of each candidate SEN-CRE, with an average spacing of 54.5□±□18.3 (mean ± SD) base pairs per sgRNA. The lentiviral sgRNA libraries targeting 9p21.3 SEN-CREs or non-targeting control (NTC) sgRNAs were transduced into MSCs stably expressing KRAB-dCas9-MeCP2 **(Fig. 2A)**. A replicative senescence model was applied to age the MSCs while maintaining ∼2,000x cell-sgRNA coverage during each passaging to ensure library representation. The NTC group served as a reference for the timing of senescence, allowing determination of whether pooled inhibition of 9p21.3 SEN-CREs could delay senescence **(Fig. 2A)**. A divergence in growth curves between two groups was observed from day 34 (passage 6; P6) to day 62 (P9), when cells in the NTC group ceased growth. At day 43 (P7) **(Fig. 2B)**, the 9p21.3 SEN-CRE group displayed more Ki67-positive proliferating cells compared to NTC group **(Fig. 2C)**, indicating that inhibition of certain 9p21.3 SEN-CREs alleviates cellular senescence in MSCs.

**Figure 2.**
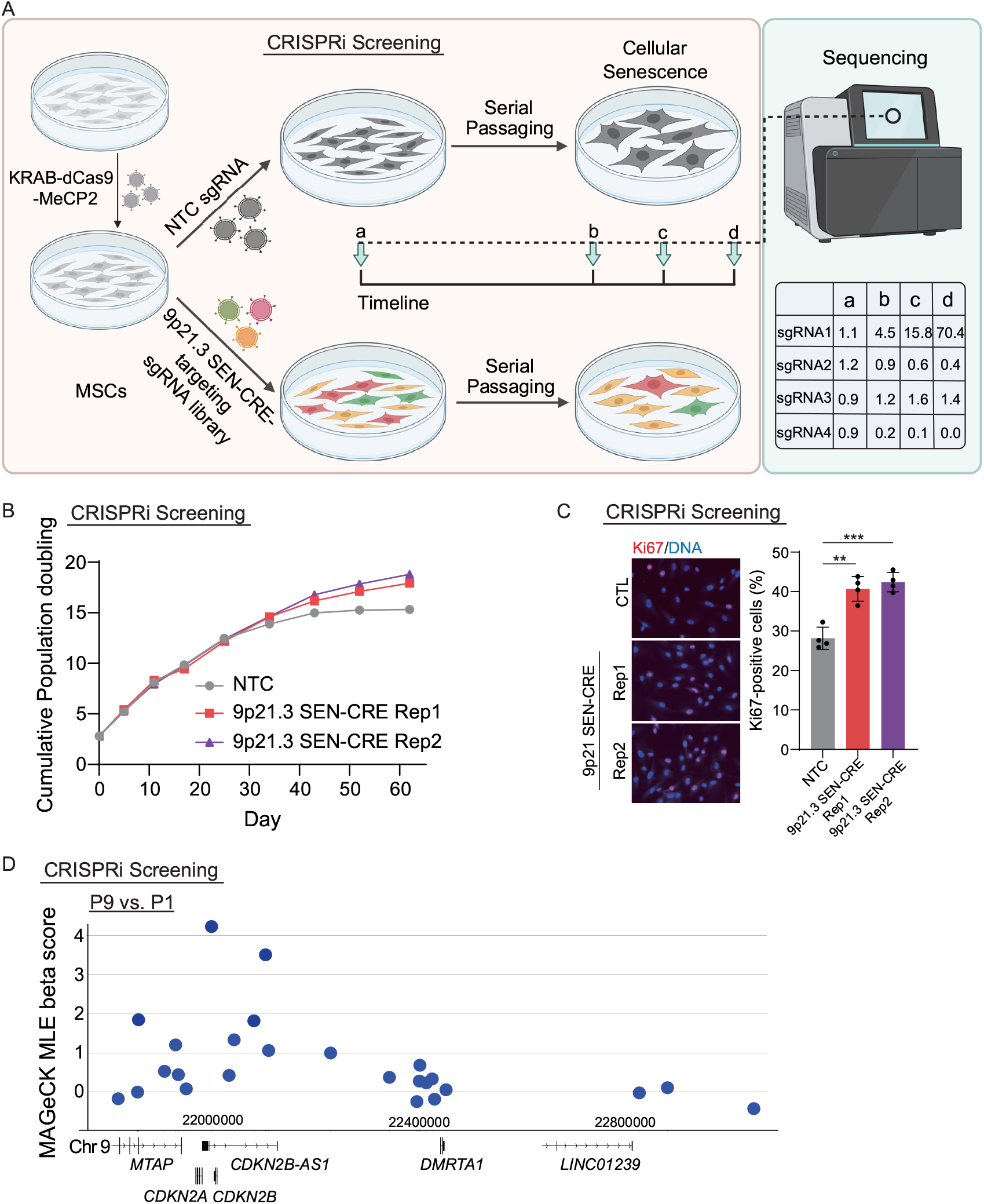
CRISPRi screen reveals 9p21.3 SEN-CREs capable of decelerating senescence. **(A)** Schematic of the CRISPR interference (CRISPRi) screen. MSCs expressing KRAB-dCas9-MeCP2 were transduced with the non-targeting control (NTC) sgRNA or the sgRNA library targeting 9p21.3 SEN-CREs. Cells were serially passaged to induce senescence and sgRNA representation was measured by sequencing. **(B)** Cumulative population doubling curve for MSCs transduced with KRAB-dCas9-MeCP2 and either the NTC or the 9p21.3 SEN-CRE sgRNA library. **(C)** Immunofluorescence staining for the proliferation marker Ki67 (red) in control (NTC) and pooled 9p21.3 SEN-CREs-inhibited MSCs at day 43 (P7). Data were presented as mean±SD. n = 3 independent images with more than 500 nuclei. ***p < 0.001. **(D)** MAGeCK MLE analysis of sgRNA enrichment at late passage (P9) versus early passage (P1). The beta-score for each SEN-CRE is plotted according to its genomic position on chromosome 9. A beta-score > 0 indicates a positive correlation with the phenotype, meaning that inhibition of SEN-CRE delayed senescence.

Next, we performed sequencing-based sgRNA enrichment analysis from the starting population (P1) through endpoint populations (P6 to P9). While sgRNA representation was well maintained at P1, diversity increased in later passages **(Fig. S2A)**, accompanied by a gradual decrease in the number of winner sgRNAs (defined as having a normalized count > 1) **(Fig. S2B)**, suggesting that senescence-resistant cells expressing winner sgRNAs became increasingly dominant during serial passaging.

We then conducted CRE-based analysis using the MAGeCK-MLE algorithm^25^, comparing sgRNA counts in P9 versus P1 cells from the 9p21.3 SEN-CRE group. As anticipated, inhibition of most SEN-CREs was correlated with delayed senescence, as reflected by positive beta scores (β > 0) **(Fig. 2D)**. Notably, the SEN-CREs with relatively stronger impact on delaying senescence were primarily enriched in regions close to *MTAP, CDKN2A, CDKN2B* and *CDKN2B-AS1*, whereas CREs near *DMRTA1* and *LINC01239* showed minimal effect **(Fig. 2D)**.

### CRISPRa screening reveals 9p21.3 SEN-CREs capable of accelerating senescence

To assess whether activation of 9p21.3 Sen-CREs can accelerate senescence, we performed a short-term CRISPR activation (CRISPRa) screen in MSCs stably expressing dCas9-VP64 using the same sgRNA libraries targeting 9p21.3 SEN-CREs **(Fig. 3A)**. We assessed senescence by measuring senescence-associated β-galactosidase (SA-β-gal) activity using SPiDER-βGal^26^. MSCs with relatively high or low SA-β-gal activities were separated and collected for sequencing **(Fig. 3B)**.

**Figure 3.**
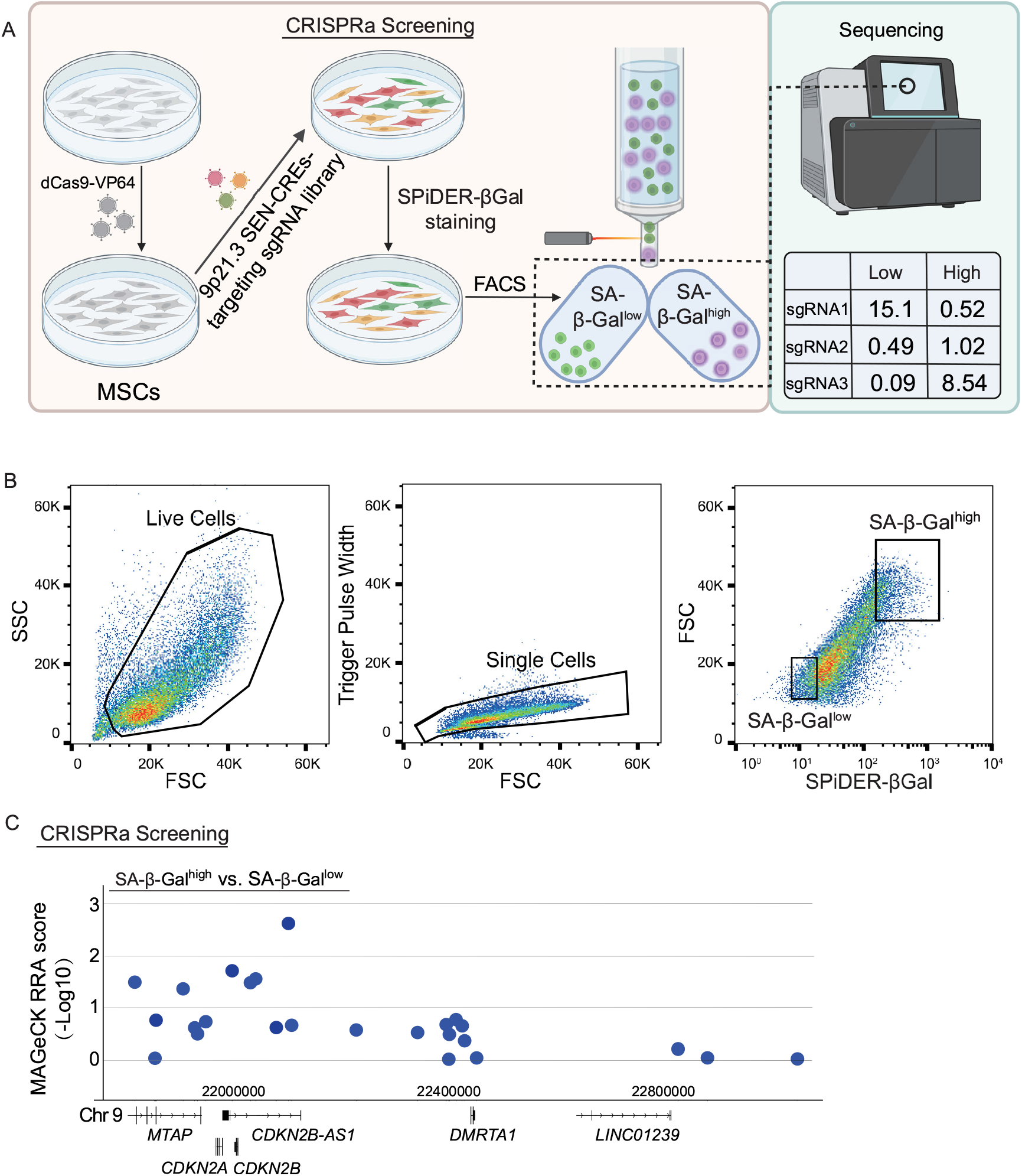
CRISPRa screening reveals 9p21.3 SEN-CREs capable of accelerating senescence. **(A)** Schematic of the CRISPR activation (CRISPRa) screen. MSCs expressing dCas9-VP64 were transduced with the 9p21.3 SEN-CRE sgRNA library. After puromycin selection, cells were stained for senescence-associated β-galactosidase (SA-β-gal) and sorted by FACS into SA-β-gal^low^ and SA-β-gal^high^ populations for sgRNA sequencing. **(B)** Representative FACS plots showing the gating strategy for live, single cells and the separation of SA-β-gal^low^ and SA-β-gal^high^ populations. **(C)** MAGeCK analysis showing SEN-CREs relatively enriched in the SA-β-gal^high^ population compared to the SA-β-gal^low^ population. The RRA score for each SEN-CRE is plotted against its genomic position, highlighting CREs whose activation accelerates senescence.

The sgRNA enrichment analysis revealed that sgRNAs targeting specific SEN-CREs were depleted in the SA-β-gal^low^ population **(Fig. S3A)**, suggesting that activation of these sgRNA-targeted CREs rapidly induces senescence. Consistent with the CRISPRi results, CRE-based analysis, comparing sgRNA counts between SA-β-gal^low^ and SA-β-gal^high^ populations, showed the most significant SEN-CREs accelerating senescence upon activation were close to *MTAP, CDKN2A, CDKN2B* and *CDKN2B-AS1* **(Fig. 3C)**.

### Identification of SenReg-CREs with bidirectional effects on senescence

To identify senescence-regulating CREs (SenReg-CREs) with bidirectional effects—accelerating senescence upon activation while delaying it upon inhibition—we cross-analyzed rankings from complementary CRISPRa and CRISPRi screens **(Fig. 4A)**. A CRE in the *CDKN2A/CDKN2B* intergenic region (named iSenReg-CRE; ranked 1st in CRISPRi and 2nd in CRISPRa) and a CRE distal from *CDKN2A/CDKN2B* (named dSenReg-CRE; ranked 2nd in CRISPRi and 1st in CRISPRa), emerged as the top candidates **(Fig. 4A-4B)**.

**Figure 4.**
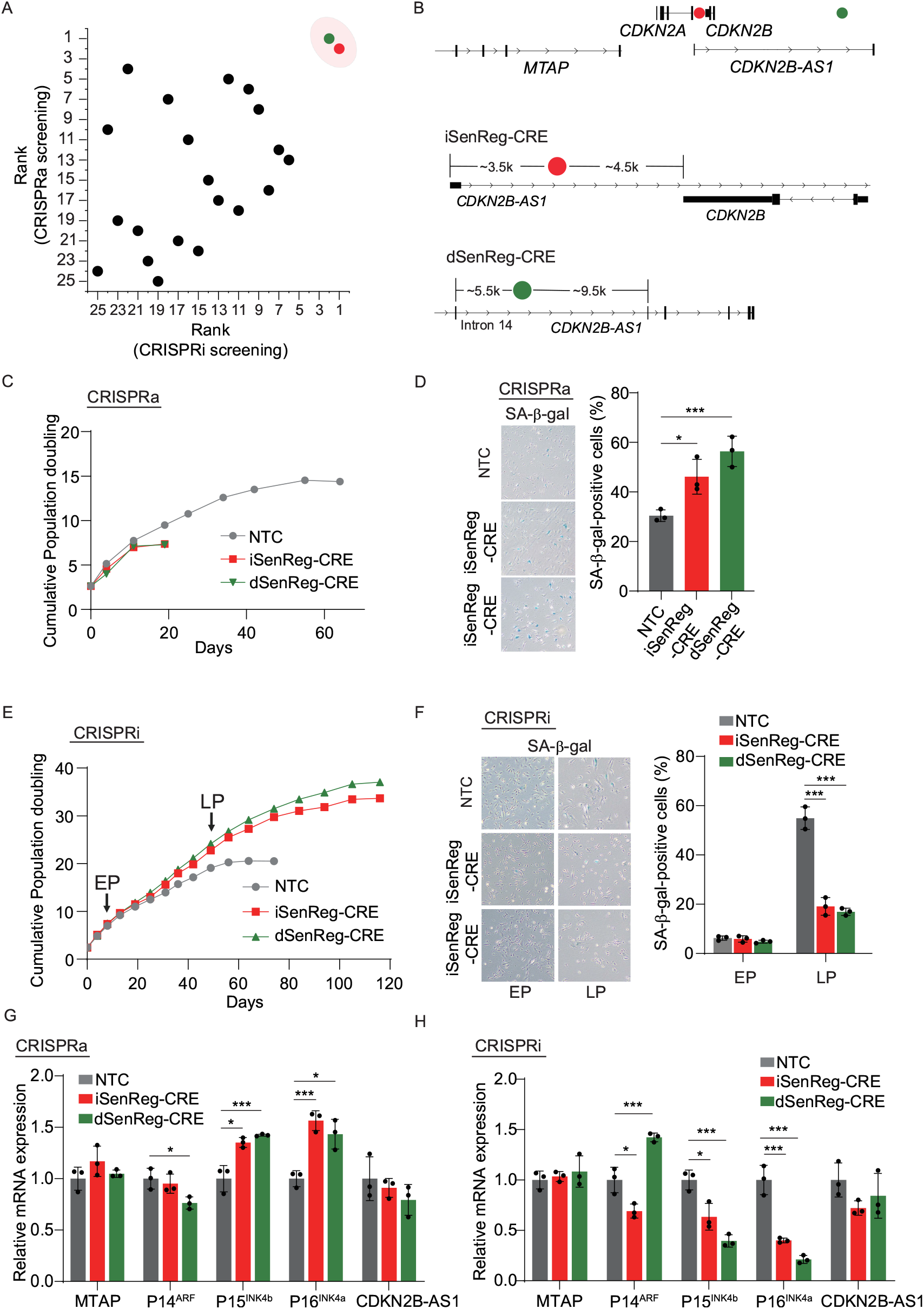
Identification of characterization of senescence-regulating CREs (SenReg-CREs). **(A)** Dot plot showing the rank of each SEN-CREs in the CRISPRa (y-axis) and CRISPRi (x-axis) screens. The selected SenReg-CREs with bidirectional effects are highlighted. **(B)** Genomic location of intergenic SenReg-CRE (red) and distal SenReg-CRE (green) relative to their nearest exons. **(C)** Cumulative population doublings for CRISPRa-MSCs expressing sgRNAs targeting NTC, iSenReg-CRE, or dSenReg-CRE. **(D)** SA-β-gal staining and quantification in CRISPRa-MSCs at P3 (3 passage after selection of sgRNA-expressing CRISPRa-MSCs). Data were presented as mean±SD. n = 3 biological replicates, *p <0.01, ***p < 0.001. **(E)** Cumulative population doublings for CRISPRi-MSCs expressing sgRNAs targeting NTC, iSenReg-CRE, or dSenReg-CRE. EP, Early Passage, P3, 3 passage after selection of sgRNA-expressing CRISPRi-MSCs; LP, Late Passage, P10, 10 passage after selection of sgRNA-expressing CRISPRi-MSCs. **(F)** SA-β-gal staining and quantification in CRISPRi-MSCs at EP and LP. Repression of iSenReg-CRE reduces the percentage of senescent cells. Data were presented as mean±SD. n = 3 independent images with more than 500 nuclei, ***p < 0.001. **(G)** Relative mRNA expression of senescence-associated genes in CRISPRa-MSCs at P3. Data were presented as mean± SD. n = 3 biological replicates, *p < 0.05, ***p < 0.001. **(H)** Relative mRNA expression of senescence-associated genes in CRISPRi-MSCs at P6. Data were presented as mean±SD. n = 3 biological replicates, *p < 0.05, ***p < 0.001.

### Functional validation of SenReg-CREs in regulating senescence

The bidirectional effects of these two SenReg-CREs on senescence were further validated independently. For each SenReg-CRE, a representative sgRNA was selected based on normalized counts from CRISPRi and CRISPRa screens **(Fig. S4A-S4B)**. Activation of SenReg-CREs reduced clonal formation capability **(Fig. S4A)** and accelerated replicative senescence **(Fig. 4C)**, as evidenced by an increased proportion of SA-β-gal positive cells **(Fig. 4D)** and decreased Ki67-positive proliferating cells **(Fig. S4D)** compared to NTC. In contrast, inhibition of SenReg-CREs delayed MSC senescence, leading to approximately 1.5-fold higher cumulative population doubling at the endpoint compared to NTC **(Fig. 4E)**, along with reduced SA-β-gal activity **(Fig. 4F)** and increased Ki67-positivity **(Fig. S4E)** at late passage. Importantly, MSCs remained capable of undergoing senescence at final passage, confirming that the cells were not immortalized **(Fig. S4F)**.

### SenReg-CREs control expression of P16^INK4a^ and P15^INK4b^

Given the potential distal *cis*-regulatory role of SenReg-CREs, we examined the expression of 9p21.3genes altered during MSC senescence, including *MTAP, CDKN2A* (P14^ARF^ and P16^INK4a^), *CDKN2B* (P15^INK4b^), and the long-non-coding RNA *CDKN2B-AS1*. Activation of both SenReg-CREs upregulated senescence-associated transcripts P16^INK4a^ and P15^INK4b^, whereas their inhibition downregulated these two genes **(Fig. 4E-4F)**. *MTAP* and *CDKN2B-AS1* expression remained unchanged **(Fig. 4E-4F)**. P14^ARF^ exhibited inconsistent changes between iSenReg-CRE and dSenReg-CRE modulation, suggesting it has a moderate impact on senescence. These results demonstrate that targeted inhibition of SenReg-CREs controlling P16^INK4a^ and P15^INK4b^ can effectively delay cellular senescence in MSC, highlighting a potential strategy to epigenetically modulate cellular aging.

### dSenReg-CRE contains a primate-specific SINE:MIR3

The dSenReg-CRE, which exhibited the strongest effect on senescence, is located distal from *CDKN2A* and *CDKN2B* **(Fig. 4E-4B)**. Among the 6 winner sgRNAs targeting dSenReg-CRE, 5 were enriched in its rear region **(Fig. 5A)**, which contains a short interspersed nuclear element (SINE) known as MIR3 (SINE:MIR3) **(Fig. 5B)**, a member of the mammalian-wide Interspersed repeat (MIRs) family^27^. MIR elements have been linked to enhancer^28^ or insulator^29^ activity, though direct experimental evidence is still lacking.

**Figure 5.**
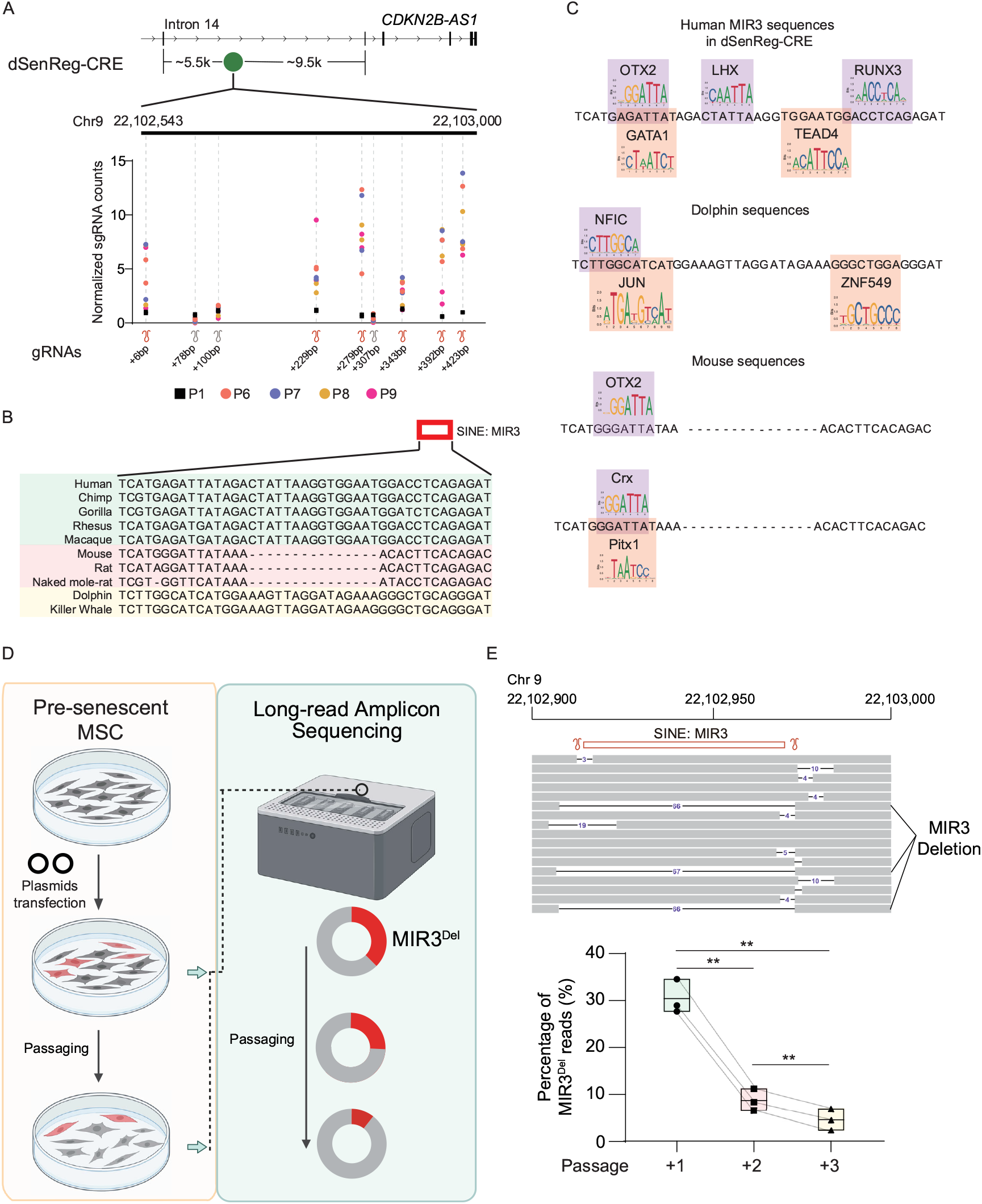
Deletion of the SINE:MIR3 element within the dSenReg-CRE accelerates MSC senescence. **(A)** Normalized sgRNA counts from the CRISPRi screen across the dSenReg-CRE locus at different passages (P1-P9). Enrichment of sgRNAs targeting the rear region at end point CRISPRi-MSCs. **(B)** Multiple sequence alignment of the regions overlapping human SINE:MIR3 element within dSenReg-CRE across various mammalian species. **(C)** Predicted transcription factor binding motifs within the human, dolphin, and mouse sequences using JASPAR human TF prediction (rows 1–3), and within the mouse sequence using JASPAR mouse TF prediction (row 4). **(D)** Schematic of the experiment determining the role of the human SINE:MIR3 in cellular senescence. Pre-senescent MSCs were transfected with plasmids expressing Cas9 and sgRNAs targeting the MIR3 flanking regions to delete the element, and the proportion of deleted alleles was monitored across subsequent passages using long-read amplicon sequencing. Depletion of MIR3-deleted reads indicates MIR3 deletion accelerates senescence, whereas enrichment of MIR3-deleted reads indicates MIR3 deletion delays senescence. **(E)** Example visualization of reads at dSenReg-CREs in the IGV browser (top). The percentage of MIR3-deleted reads decreased over three passages (bottom), Data are shown as mean ± range (minimum to maximum). n = 3 biological replicates. **p<0.01.

The SINE:MIR3 within dSenReg-CRE is highly conserved in humans and non-human primates **(Fig. 5B)**. Transcription factor (TF) binding motif analysis of the primate-specific MIR3 revealed potential interactions with multiple TFs, including OTX2, GATA1, LHX, TEAD4 and RUNX3 **(Fig. 5C)**. Cetacean-specific sequences, although the same length as the primate-specific MIR3, exhibit substantially different TF binding profiles, losing the ability to bind LHX, TEAD4, and RUNX3. Rodent-specific sequences showed poor conservation with MIR3, particularly in the central region, and retain only the binding potential for homeodomain factors such as Crx **(Fig. 5C)**. These species-specific differences in TF binding profiles suggest that primates may have evolved a distinct mechanism for epigenetic regulation of *CDKN2A/CDKN2B* and the subsequent control of senescence.

### SINE:MIR3 deletion accelerates MSC senescence

To investigate the functional role of SINE:MIR3 within dSenReg-CRE in modulating cellular senescence, we deleted SINE:MIR3 in the genome of pre-senescent MSCs using CRISPR-Cas9 with paired sgRNAs flanking the SINE:MIR3 **(Fig. 5D)**. The presence and proportion of SINE:MIR3-deleted alleles in pre-senescent MSCs were monitored continuously using long-read sequencing **(Fig. 5E)**. During serial passaging, the proportion of SINE:MIR3-deleted alleles progressively declined **(Fig. 5E)**, indicating the loss of SINE:MIR3-deleted cells from the cell population. This observation suggests that SINE:MIR3 deletion accelerates MSC senescence and that SINE:MIR3 functions as an insulator, potentially restraining *CDKN2A/CDKN2B* activation but fails during cellular senescence.

## Discussion

CRISPR screens have emerged as a powerful method enabling systematic interrogation of genetic components across diverse applications^30^. Several studies have successfully employed phenotypic CRISPR screens to study cellular senescence^31-35^ and identified novel genes as drivers of cellular aging, however, all have focused on protein-coding genes. Non-coding regions, which comprise over 98% of the human genome and contain numerous regulatory elements critical for gene expression, remain largely unexplored^36^ in the context of cell aging. This gap is particularly pronounced for complex genomic loci such as 9p21.3 locus where extensive gene deserts contain multiple regulatory elements with unclear functions. Our complementary CRISPRa and CRISPRi screens of regulatory elements across the 1.25 Mb 9p21.3 locus therefore represent a comprehensive approach to dissecting functional non-coding CREs that modulate cellular senescence through key senescence-associated genes, P16^INK4a^ and P15^INK4b^.

A particularly intriguing finding is the identification of a primate-specific SINE:MIR3 element embedded within the most potent dSenReg-CRE. MIR3 (chr9: 22102915-22102957, hg38) is located at the boundary of a strong active enhancer named ECAD5 (chr9: 22102620-22105003, hg38) in a previous study^37^. Given that the SINE:MIR3-residing region becomes accessible during cellular senescence and that MIR3 deletion accelerates senescence, MIR3 may function as a cis-regulatory insulator likely by forming boundary complexes that prevent the adjacent enhancer from activating promoters^38^, however, this insulation fails during senescence. In addition, the primate-specific SINE:MIR3 uniquely harbors binding sites for LHX, TEAD4, and RUNX3, highlighting its evolutionary specificity and potentially primate-specific mechanisms in the regulation of cellular senescence.

Recent advances in genetic enhancement in MSCs have demonstrated that targeting conserved aging pathways or genes can improve stress resistance and therapeutic potential. For example, constitutive activation of the antioxidative response gene *NRF2*^*21*^ or the longevity-associated gene *FOXO3*^*22*^ through introducing coding mutations has been reported to delay MSC senescence, enhance tissue regeneration in mice, and counteract systemic aging in monkeys^19^. However, altering coding sequences raises concerns about disrupting evolutionarily optimized mechanisms, such as the dynamic turnover of NRF2 or the phosphorylation-dependent subcellular shuttling of FOXO3. As an alternative, epigenetic manipulation of senescence-associated genes via non-coding CREs offers a promising strategy to enhance cellular function without permanently altering protein-coding sequences. A similar concept has been successfully applied clinically, as demonstrated by the first FDA-approved CRISPR-based therapy Casgevy^39^, where disruption of an erythroid-specific enhancer in the *BCL11A* gene^40^ reactivates fetal hemoglobin expression to treat sickle cell disease. We demonstrate that modulating *CDKN2A/CDKN2B* through functional CREs can effectively control cellular senescence and thus the SenReg-CREs identified in this study serve as promising targets for epigenetic modulation of cellular senescence.

We acknowledge the limitations of this study. The regulatory mechanisms we identified were characterized specifically in MSCs, and their activity and functional relevance may vary across different cell types and tissues. Therefore, validation of these SenReg-CREs in additional cellular contexts and disease models will be important to fully understand their roles in cellular senescence.

## Figure Legends

**Figure S1.**
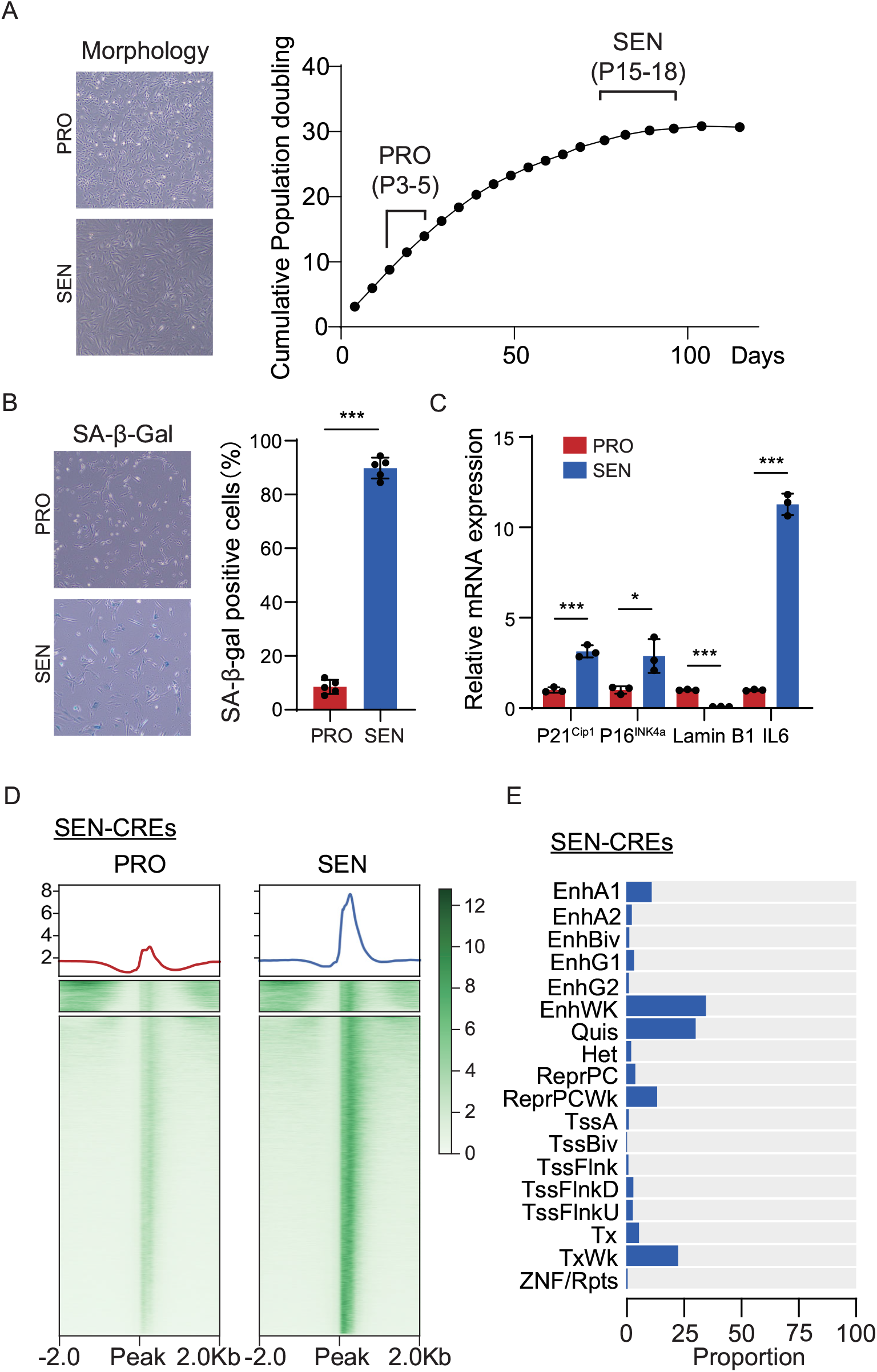
Identification of SEN-CREs during replicative senescence in MSCs. **(A)** Left: Morphology of proliferating (PRO) and senescent (SEN) MSCs. Right: Growth curve showing cumulative population doublings. **(B)** SA-β-gal staining and quantification confirming cellular senescence at late passages. ***p < 0.001. **(C)** Relative mRNA expression of senescence markers *P21*^*Cip1*^, *P16*^*INK4a*^, *Lamin B1*, and *IL6*. N = 3, *p < 0.05, ***p < 0.001. **(D)** Chromatin accessibility centered on SEN-CREs in PRO and SEN cells. **(E)** Proportion of SEN-CREs overlapping different ChromHMM chromatin states.

**Figure S2.**
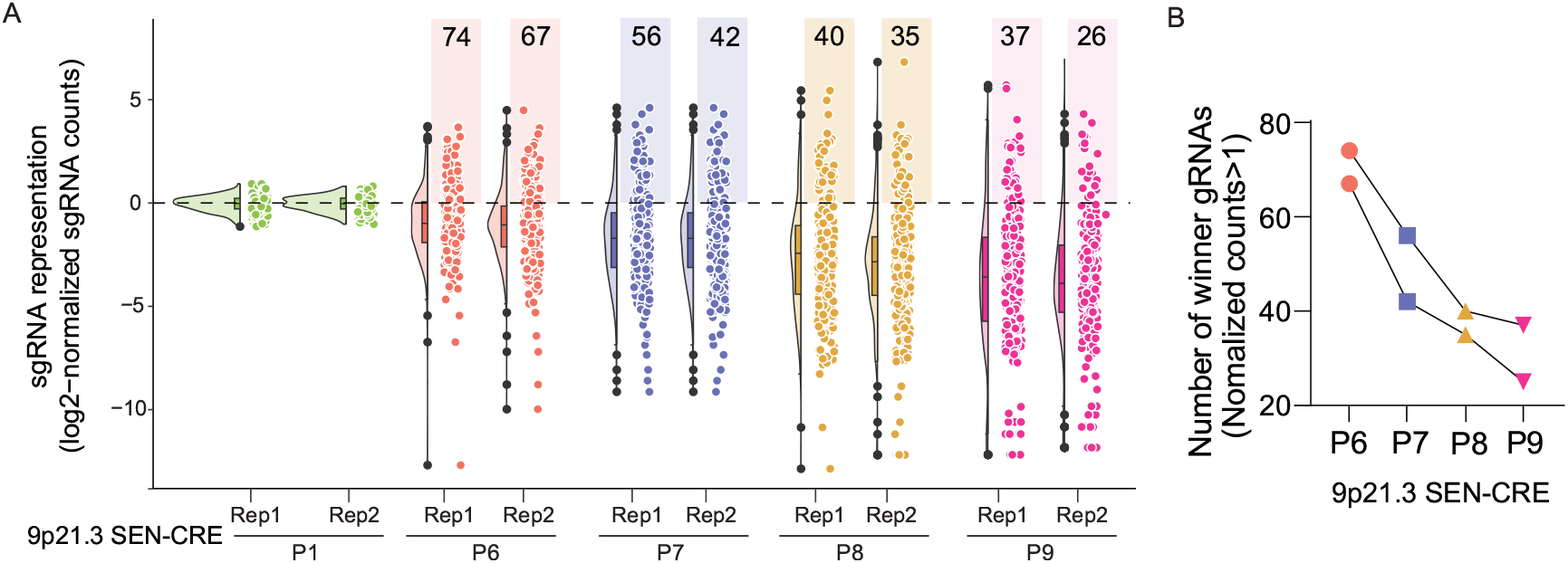
sgRNA representation analysis in CRISPRi screening. **(A)** Violin plots illustrating the distribution of log2-normalized sgRNA counts for two replicates of CRISPRi-MSCs at passages 1, 6, 7, 8, and 9. The numbers at the top indicate the number of winner sgRNAs (normalized count > 1) at each time point. **(B)** Changes in the number of winner sgRNAs from P6 to P9 in CRISPRi-MSCs.

**Figure S3.**
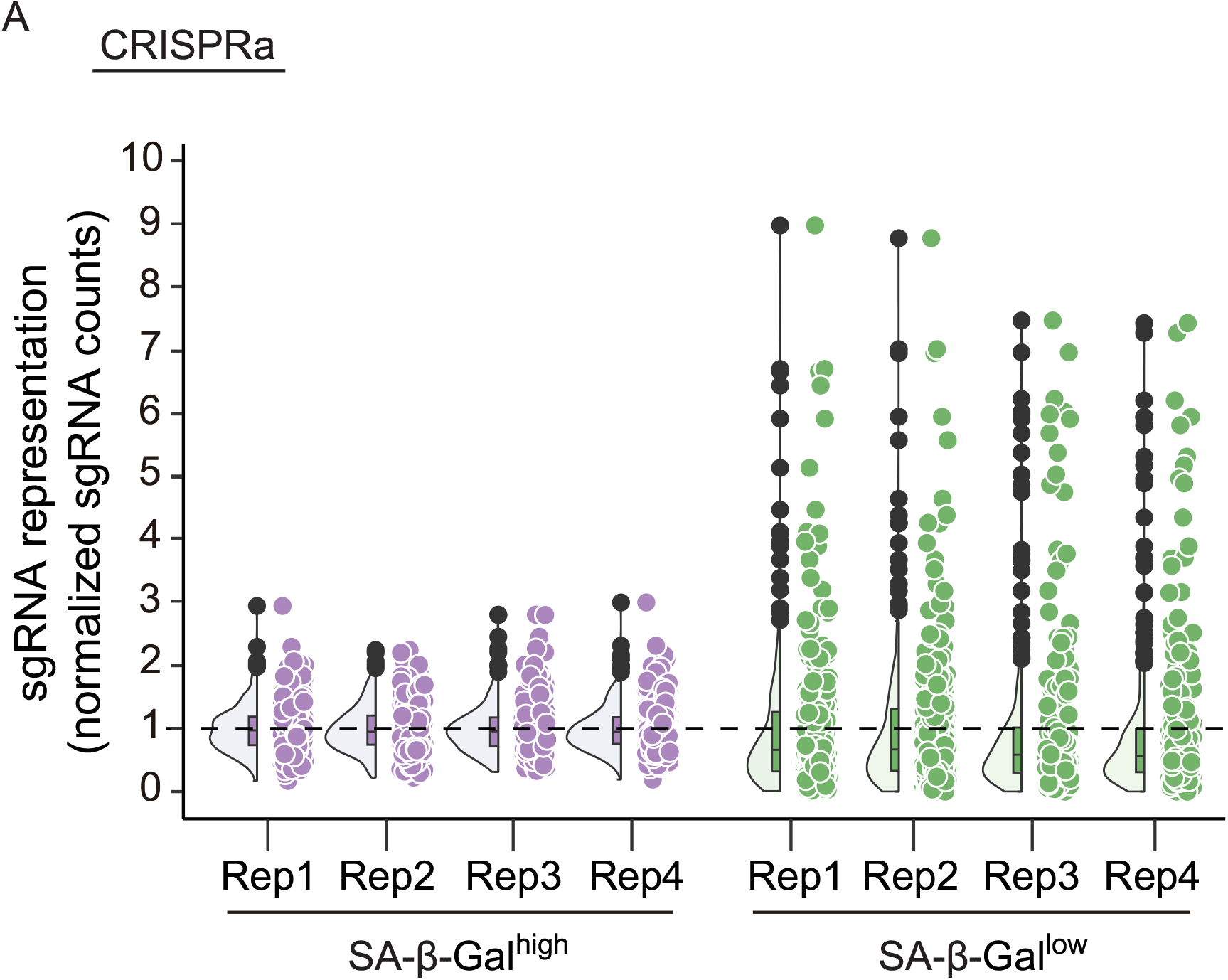
sgRNA representation analysis in CRISPRa screening. **(A)** Violin plots showing the distribution of normalized sgRNA counts from four biological replicates in the SA-β-gal^high^ (purple) and SA-β-gal^low^ (green) sorted populations.

**Figure S4.**
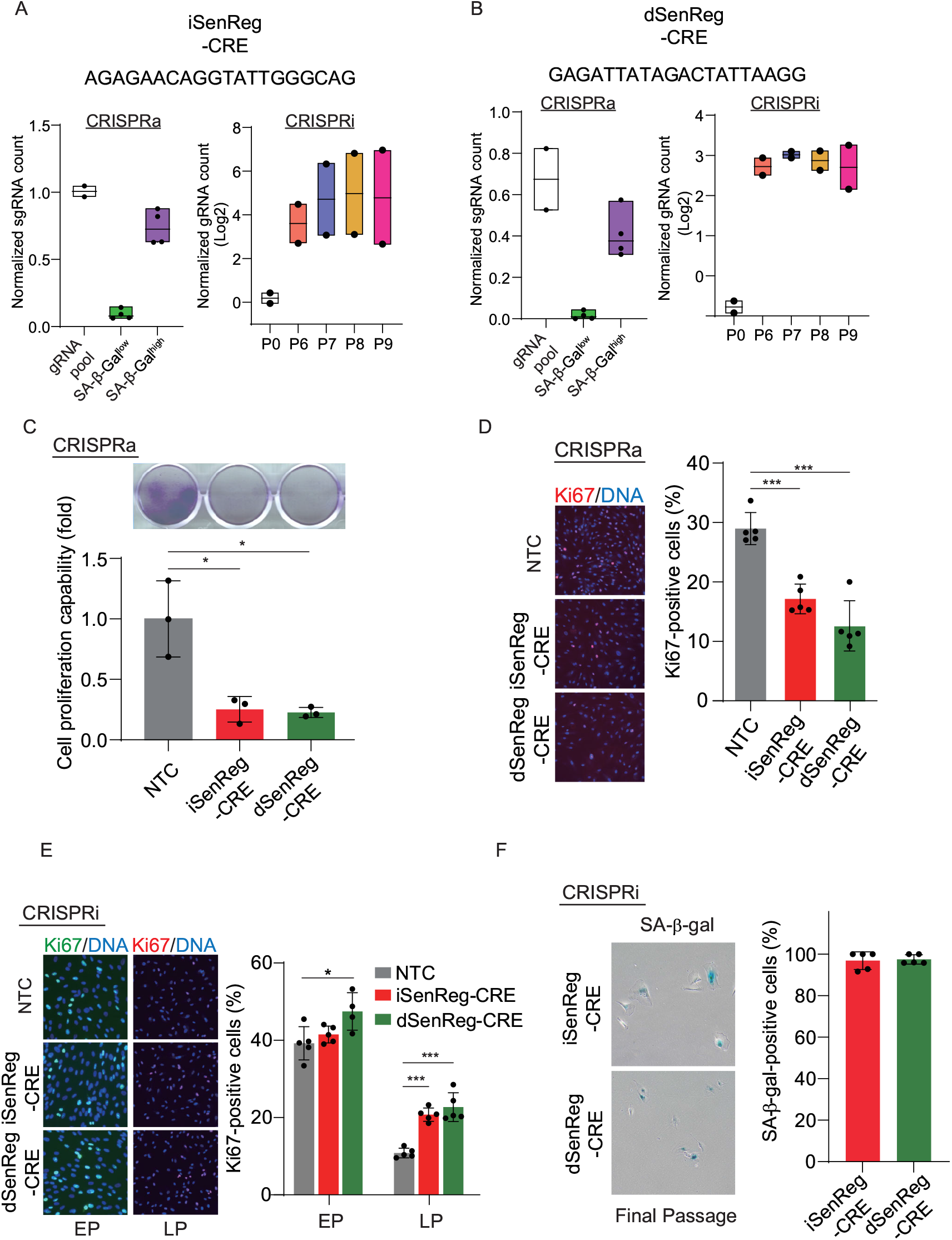
Characterization of iSenReg-CRE and dSenReg-CRE. **(A)** Box plot showing the counts of the representative sgRNA for iSenReg-CRE in the CRISPRa and CRISPRi screen. Data are shown as mean ± range (minimum to maximum). **(B)** Box plot showing the counts of the representative sgRNA for dSenReg-CRE in the CRISPRa and CRISPRi screen. Data are shown as mean ± range (minimum to maximum). **(C)** Clonal formation capability measured by crystal violet staining in CRISPRa-MSCs. n = 3 biological replicates, *p < 0.05. **(D)** Representative images and quantification of Ki67-positive cells in the CRISPRa-MSCs at P3. n = 5 independent images with more than 500 nuclei, ***p < 0.001. **(E)** Representative images and quantification of Ki67-positive cells in the CRISPRi-MSCs at EP and LP. n =4 or 5 independent images with more than 500 nuclei, *p < 0.05, ***p < 0.001. **(F)** Representative images and quantification of SA-β-gal staining at the final passage of the CRISPRi experiment, n = 5 independent images with more than 100 nuclei.

## Materials and Methods

### Cell culture

Human H7 ESCs (WiCell Research) were maintained on mitomycin C-inactivated mouse embryonic fibroblast (MEF) feeder (Thermo Fisher) in human ESC medium (DMEM/F12 (Invitrogen), 20% Knockout Serum Replacement (Invitrogen), 0.1 mM non-essential amino acids (NEAA, Invitrogen), 2 mM GlutaMAX (Invitrogen), 55 μM β-mercaptoethanol (Invitrogen), and 10 ng/ml FGF2 (Thermo Fisher)) and were differentiated into human MSC as previously described. Human MSCs were cultured on Gelatin-coated plate in MSC culture medium (MEMa (Gibco), 10% fetal bovine serum (Hyclone) and 1 ng/mL bFGF (Thermo Fisher)).

All cell culture media were supplemented with 1% penicillin/streptomycin (Gibco) and 50ug/mL Normocin (InvivoGen) to prevent cells from microbial contaminations. Cultured cells are routinely tested for mycoplasma contamination using Mycoplasma PCR Detection Kit (ABM).

### Mesenchymal stromal cell differentiation

hMSCs were differentiated from hESCs as previously described with minor modification. Briefly, embryoid bodies (EB) were first generated using ESCs cultured on MEF feeder and cultured on ultra-low attachment plates (Corning) using EB medium (human ESC medium with 4 ng/ml FGF2) for three days, the EBs were then plated on Matrigel-coated plates in hMSC differentiation medium (αMEM (Gibco), 10% fetal bovine serum (GeminiBio), 1% penicillin/streptomycin (Gibco), 10 ng/mL FGF2 (Thermo Fisher) and 5 ng/mL TGFβ (Thermo Fisher)) for around 10 days till fibroblast-like cells were confluent. These fibroblast-like cells were maintained in hMSC culture medium on Gelatin-coated 10cm dishes for two passages and were further sorted by the BD Influx cell sorter to purify CD73/CD90/CD105 tri-positive MSCs.

### Replicative senescence assay

Cell population doubling was calculated as previously described. Briefly, MSCs were serially passaged and the number of cells was counted. Population doubling per passage was calculated as log2 (number of cells harvested/number of cells seeded). Cumulative population doublings of the cells were calculated and plotted to days. Cells were considered to have entered senescence when the number of cells harvested is lower than the number initially seeded.

### RT-qPCR

Cells were pelleted and total RNA was extracted using RNeasy Mini Plus Kit (Qiagen). Then HiScript IV RT SuperMix for qPCR (+gDNA wiper) (Vazyme) was used to eliminate genomic DNA and generate cDNA. RT-qPCR was performed with PowerUp SYBR Green Master Mix (ThermoFisher) in QuantStudio 6 Pro Real-Time PCR System (ThermoFisher). All primer sequences for qPCR are listed below: P14^ARF^-Forward: 5′-GGGTTTTCGTGGTTCACATCC-3′, Reverse: 5′-CTAGACGCTGGCTCCTCAGTA-3′; P15^INK4B^-Forward: 5′-CGCTGCCCATCATCATGAC-3′, Reverse: 5′-CTAGTGGAGAAGGTGCGACA-3′; P16^INK4A^-Forward: 5′-ATGGAGCCTTCGGCTGACT-3′, Reverse: 5′-GTAACTATTCGGTGCGTTGGG-3′; P21-Forward: 5′-CGATGGAACTTCGACTTTGTCA-3′, Reverse: 5′-GCACAAGGGTACAAGACAGTG-3′; CDKN2B-AS1-Forward: 5′-ACACATCAAAGGAGAATTTTCTTGG-3′, Reverse: 5′-GTACTGACTCGGGAAAGGATTC-3′; MTAP-Forward: 5′-ACCACCGCCGTGAAGATTG-3′, Reverse: 5′-GCATCAGATGGCTTGCCAA-3′; Lamin B1-Forward: 5′-GTAAGCACTGATTTCCATGTCCA-3′, Reverse: 5′-GAAAAAGACAACTCTCGTCGCA-3′; IL6-Forward: 5′-ACTCACCTCTTCAGAACGAATTG-3′, Reverse: 5′-CCATCTTTGGAAGGTTCAGGTTG-3′.

### RNA-seq analysis

RNA-seq data of human embryonic stem cell-differentiated MSC were obtained from NCBI Gene Expression Omnibus (GEO; accession number GSE84694). Proliferating MSC samples correspond to GSM2247746 and GSM2247747, and senescent MSC samples to GSM2247750 and GSM2247751. Adapter trimming and quality filtering were performed using Trim Galore (v0.6.10). Cleaned reads were aligned to the human reference genome (hg38) using STAR (2.7.11b) with the parameter--outFilterMismatchNoverLmax 0.05 to limit mismatches relative to read length. BAM files were converted to bigWig format using bamCoverage from deepTools (v3.5.6), normalizing using RPKM. The resulting tracks were visualized in Integrative Genomics Viewer (IGV).

### ATAC-seq library preparation

ATAC-seq of PRO (P3) and SEN (P17 and P18) were performed using Omni-ATAC-seq protocol as previously described (ref) with minor modifications. Briefly, 50,000 cells per sample were collected, washed with PBS, and lysed in cold lysis buffer (10 mM Tris-HCl pH 7.4, 10 mM NaCl, 3 mM MgCl_2_) containing 0.1% NP-40, 0.1% Tween-20, and 0.01% digitonin for 3 minutes on ice. Nuclei were then washed with cold lysis buffer containing 0.1% Tween-20, and immediately subjected to transposition. The transposition reaction was carried out using 2.5 μl Tn5 transposase (Illumina), 25 μl 2× TD buffer, 16.5 μl PBS and 5 μl nuclease-free water in a total volume of 50 μl, incubated at 37°C for 30 minutes with gentle mixing (1,000 rpm). Following tagmentation, DNA was purified using a DNA Clean & Concentrator Kit (Zymo). Library preparation was performed using NEBNext Q5 Hot Start HiFi PCR Master Mix with the number of cycles determined empirically by qPCR to avoid overamplification. Final libraries were purified with AMPure XP beads (Beckman Coulter) using a double-sided selection (0.5-1.8x) and assessed for size distribution and concentration using an Agilent Bioanalyzer and Qubit fluorometer. The ATAC-seq libraries were sequenced using Illumina NextSeq 500 (Pair-end 75bp) at Center for Epigenomics and Computational Genomics Core at Albert Einstein College of Medicine Genomic center.

### ATAC-seq analysis

ATAC-seq data were processed following ENCODE ATAC-seq pipeline (https://github.com/ENCODE-DCC/atac-seq-pipeline) with minor modification. Briefly, raw paired-end FASTQ files were cleaned using Trim Galore (v0.6.10) and were aligned to the human reference genome (hg38) using Bowtie2 (v2.2.6). PCR Duplicate reads were removed using MarkDuplicates from Picard (v1.1256) and mitochondrial reads were removed by filtering out reads mapped to chromosome chrM. To correct for Tn5 insertion bias, aligned BAM files were further processed using alignmentSieve (deepTools v3.5.6) with the --ATACshift option. Normalized ATAC-seq signal tracks were created from shifted BAM files using bamCoverage (deepTools v3.5.6). Reads were normalized to RPGC using the human genome size. Output files in bigWig format were generated for visualization and analysis. Peak calling was performed using Genrich (v0.6.1) on non-shifted BAM files, with the ATAC-seq mode (option -j) to enable ATAC-seq specific Tn5 insertion site shifting. Senescence-specific ATAC-seq peaks (defined as SEN-CREs) were identified by subtracting peaks found in the PRO MSC from those in SEN MSC using bedtools subtract with the -A option. The computeMatrix and plotHeatmap (deepTools v3.5.6) were used to visualize ATAC-seq signal enrichment around SEN-CREs.

### Immunofluorescence staining

Cells were fixed in 4 % paraformaldehyde at room temperature (RT) for 15 min, permeabilized in 0.4% Triton X-100/PBS at RT for 10 min. After blocking with 10% donkey serum (Jackson ImmunoResearch Labs)/PBS for 1 h, cells were incubated with Ki67-APC (Thermo Fisher, 17-5699-42) antibodies at 4 °C overnight. Nuclei were stained with Hoechst 33342 (Thermo Fisher, 62249). Images were captured using EVOS M7000 Imaging System (Thermo Fisher).

### SA-β-gal staining assay

The SA-β-gal staining of hMSCs was conducted as previously described. Briefly, cells were washed with PBS and fixed at room temperature for 2-5 minutes using a fixation buffer containing 2% formaldehyde and 0.2% glutaraldehyde. After fixation, cells were incubated overnight at 37□°C with freshly prepared staining solution. SA-β-gal-positive and total cell numbers were quantified using ImageJ software, and the percentage of SA-β-gal-positive cells was calculated for statistical analysis.

### sgRNA library design

The Python script design_library.py (https://github.com/fengzhanglab/Screening_Protocols_manuscript), as described by Joung et al Nature Protocols 2017, was used to generates sgRNAs targeting 25 SEN-CREs within 1.25Mb spanning protein-coding genes *MTAP, CDKN2A, CDKN2B, DMRTA1* and long-noncoding RNA *CDKN2B-AS1, LINC01239* at 9p21.3. These gRNAs were designed to meet specific criteria including GC%>25%, no homopolymer, and no more than 3 off-target sites. Totally, 285 sgRNAs were designed to target these SEN-CREs, the sgRNA sequences were listed in Table S1.

### sgRNA plasmid construction

For the sgRNA libraries used in CRISPR screening, sgRNAs flanked by the sequences TATATATCTTGTGGAAAGGACGAAACACCG and GTTTAAGAGCTATGCTGGAAACAGC were synthesized by Twist Bioscience. sgRNA oligo library was amplified using NEBNext Q5 Hot Start HiFi PCR Master Mix with oligo-forward primer: GTAACTTGAAAGTATTTCGATTTCTTGGCTTTATATATCTTGTGGAAAGGACGAAACACC and oligo-reverse primer: ACTTTTTCAAGTTGATAACGGACTAGCCTTATTTTAACTTGCTATTTCTAGCTCTAAAAC. The amplified oligos were purified using DNA Clean & Concentrator Kit (Zymo) and cloned into lentiGuide-Puro (Addgene, 52963) backbone cleaved by FastDigest Esp3I (Thermo Fisher, FD0454) using NEBuilder HiFi DNA Assembly (NEB). The recombinant products were purified and concentrated through isopropanol precipitation. Eluted library was electroporated using Endura ElectroCompetent cells (BiosearchTechnologies), cultured on LB plate overnight and midi-prepped using Macherey-Nagel NucleoBond Xtra Midi EF kit. The sgRNA regions of sgRNA plasmid library were amplified and sequenced using Amplicon-seq service (GENEWIZ) to determine sgRNA representation. For individual validation, representative sgRNAs targeting iSenReg-CRE (AGAGAACAGGTATTGGGCAG) and dSenReg-CRE (GAGATTATAGACTATTAAGG)were cloned into lentiGuide-Puro (Addgene, 52963) backbone cleaved by FastDigest Esp3I (Thermo Fisher, FD0454).

### CRISPR lentivirus packaging

Lentivirus plasmids including lentiCRISPRi(v2)-Blast (Addgene, 170068), lenti-dCAS-VP64_Blast (Addgene, 61425), LentiGuide-Puro targeting control scramble (GCTTAGTTACGCGTGGACGA), 9p21.3 SEN-CREs, and targeting two SenReg-CREs. Lentivirus packaging was performed in HEK293T cells by transfection of lentiviral vector plasmids together with pMD2.G (Addgene, 12259) and psPAX2 (Addgene, 12260) using Lipofectamine 3000 (Thermo Fisher). Media was replaced after overnight incubation. Lentiviral supernatant was collected 48 hours post-transfection, purified by passing through 0.22 um filters (Millipore) and concentrated using Lenti-X Concentrator (Takara).

### CRISPRi screening

MSCs at passage 3 were transduced with lentiCRISPRi(v2)-Blast lentiviruses at an MOI of 0.5 with 10 μg/ml polybrene (Millipore) to generate CRISPRi-MSCs. Transduced cells were selected with Blasticidin (5 ug/mL), expanded for 2 additional passages under Blasticidin selection, and cryopreserved for screening and individual validation studies. CRISPRi-MSCs were further transduced with either control sgRNA or sgRNA library lentiviruses targeting 9p21.3 SEN-CREs at an MOI of 0.5 with > 2,000x coverage (∼1.2 million cells per replication). Transduced cells selected with Puromycin (2 ug/mL) were used in replicative senescence assay. To maintain the complexity of the sgRNA library within the cell population, 800,000 cells per replicate (∼ 2,700× coverage) were seeded during each cell passaging. The remaining cells after passaging were harvested and stored at −80°C. For CRISPRi screening, cells at P1, P7, P8 and P9 were used for sgRNA sequencing.

### CRISPRa screening

MSCs at passage 3 were transduced with lenti-dCAS-VP64_Blast lentiviruses at an MOI of 0.5 to generate CRISPRa-MSCs. Transduced cells were selected with Blasticidin (5 ug/mL), expanded for 2 additional passages under selection, and cryopreserved for screening and individual validation studies. CRISPRa-MSCs were further transduced with either control sgRNA or sgRNA library lentiviruses targeting SEN-CREs at an MOI of 0.5 with >2,000x coverage (∼1.2 million cells per replicate). Transduced cells were selected with Puromycin (2 ug/mL) 48 hours post-transduction. Cells were expanded for one passage and stained using SPiDER-βGal Cellular Senescence Detection Kit (Dojindo, SG04) according to the manufacturer’s manual. Attached MSCs were incubated with Bafilomycin A1 working solution at 37 °C for 1 hour to inhibit endogenous β-galactosidase activity. The SPiDER-β-Gal working solution was further added to stain cells at 37 °C for 30 minutes. Next, cells were then dissociated and sorted using a SONY MA900 cell sorter to enrich for β-Gal^high^ and β-Gal^low^ populations, corresponding to the around top 10% and bottom 10% of β-Gal signal intensity, respectively. The collected cells were pelleted for sgRNA sequencing.

### Sequencing of sgRNA Libraries from CRISPR screens

Genomic DNA was extracted from harvested cells following CRISPRa or CRISPRi screening using the QIAGEN DNeasy Blood & Tissue Kit. To amplify sgRNA target regions, indexed primers were used to perform PCR on genomic DNA from each sample. PCR using NEBNext Ultra II Q5 Master Mix was carried out in triplicate with 20 cycles per reaction to minimize amplification bias and error. The resulting PCR products were purified using the DNA Clean & Concentrator-25 Kit (Zymo Research) and subsequently pooled. Pooled libraries were subjected to next-generation sequencing on an Illumina NextSeq platform at the Center for Epigenomics and Computational Genomics Core at Albert Einstein College of Medicine.

### sgRNA-based analysis

Raw sgRNA counts were quantified from demultiplexed FASTQ files using the Python script count_spacers.py (https://github.com/fengzhanglab/Screening_Protocols_manuscript), as described by Joung et al., *Nature Protocols*, 2017. The script identifies sgRNA reads by locating the 20 bp spacer sequence immediately downstream of the constant sequence “CGAAACACC” and matches them to a reference library of designed sgRNAs. Only perfect matches to the reference spacer sequences were included to ensure specificity and accuracy. Normalized sgRNA counts were calculated using the formula: (raw count of each sgRNA / total counts in the sample) × total number of sgRNAs in the library, to account for differences in sequencing depth across samples.

### CRE-based analysis

CRE-based analysis was performed using MAGeCK (Model-based Analysis of Genome-wide CRISPR-Cas9 Knockout). For the CRISPRi screening, which involved a long-term selection comparing cells at the last passage (P9) versus the first passage (P1), the MAGeCK-MLE (maximum likelihood estimation) algorithm was used as MAGeCK-MLE is well-suited for time-course or prolonged selection experiments. For the CRISPRa screen, which compared SA--high versus bgal-low cell populations in a short-term, endpoint-based selection, the MAGeCK-RRA (robust rank aggregation) algorithm was applied. RRA ranks sgRNAs based on their relative enrichment or depletion between two conditions and is optimized for binary comparisons such as sorted populations.

### Transcription factor binding prediction

We used the JASPAR database for transcription factor (TF) binding predictions. A total of 251 ChIP-seq-based profiles were employed to predict the binding of human TFs to sequences from human, mouse, and dolphin. For predictions of mouse TFs binding to mouse sequences, all 224 mouse-specific profiles were used. The relative profile score threshold was set at 90%.

### SINE:MIR3 deletion

sgRNAs targeting the 5-prime (ATGGTCCTTGCTCTCATCTG) and 3-prime (TCTCATGAATTTAGCTAACT) of SINE:MIR3 were cloned into lentiCRISPR v2 (Addgene, 52961) backbone cleaved by FastDigest Esp3I (Thermo Fisher, FD0454). Both plasmids were transfected into pre-senescent MSCs (P12) using Lipofectamine Stem Reagent (Thermo Fisher). Transfected MSCs were selected using 1ug/mL Puromycin (InvivoGen) 2 days post-transfection for 3 days and maintained without Puromycin for another 4 days. From the next passage (P1), cells were serially passaged at 10E5 per 6 wells. Remaining cells after passaging were collected for long-read sequencing.

### Long-read amplicon sequencing of the MIR3 region

Genomic DNA was extracted using the QIAGEN DNeasy Blood & Tissue Kit. A 1,391 bp region spanning chr9:22101967–22103357 (hg38) was amplified by PCR with primers TTAGCTCCAGAGACTGCAACCAACTGCCTA and ACTTCTCAATGCCCCAATAGCCAAGCTCC, using NEBNext Ultra II Q5 Master Mix. Each sample was amplified in duplicate with 30 cycles per reaction. PCR products were purified with the DNA Clean & Concentrator-25 Kit (Zymo Research) and subsequently pooled. The pooled PCR products were sequenced using PCR-EZ service (Genewiz). FASTA files generated by Genewiz were mapped with BWA-MEM and visualized in IGV. The number of MIR3-deleted reads and total reads were recorded for statistical analysis.

## Data availability

All sequencing data have been deposited to Sequence Read Archive in the National Center for Biotechnology Information. The accession numbers were as follows: ATAC-seq of ESC-derived MSC (PRJNA1272555).

